# Natural transformation specific DprA coordinate DNA double strand break repair pathways in heavily irradiated *D. radiodurans*

**DOI:** 10.1101/2023.07.11.548530

**Authors:** Dhirendra Kumar Sharma, Ishu Soni, Hari S. Misra, Yogendra Singh Rajpurohit

## Abstract

*Deinococcus radiodurans* exhibits remarkable survival under extreme conditions, including ionizing radiation, desiccation, and various DNA-damaging agents. It employs unique repair mechanisms, such as single-strand annealing (SSA) and extended synthesis-dependent strand annealing (ESDSA), to efficiently restore damaged DNA fragments. In this study, we investigate the regulatory role of the NT-specific protein DprA in DNA repair pathways following acute gamma radiation exposure. Our findings demonstrate that the absence of DprA leads to rapid repair of gamma radiation-induced DNA double-strand breaks (DSBs), with diminished involvement of the ESDSA pathway. Furthermore, our data suggest that the SSA pathway becomes the primary mechanism for DNA DSB repair in the absence of DprA. Overall, our results highlight the regulatory function of DprA in modulating the choice between SSA and ESDSA pathways for DNA repair in the radiation-resistant bacterium *D. radioduransx*.

## Introduction

Natural transformation (NT) is a regulated form of horizontal gene transfer in bacteria, serving various purposes such as genome evolution, DNA repair, and fulfilling nutritional needs [1]. The process involves interactions between extracellular DNA (eDNA) and pili structures, followed by the movement of eDNA into the cytoplasm with the assistance of specific proteins [2]. To protect the translocated DNA from degradation, proteins like DprA and single-stranded DNA binding proteins (SSB) are crucial. DprA, as a member of the natural transformation-specific recombination mediator proteins (RMP) family, facilitates the loading of RecA on single-stranded DNA (ssDNA) [3, 4]. The fate of the transformed DNA can be either chromosomal integration or reconstitution into an autonomous plasmid [5]. The impact of natural transformation on bacterial evolution and genome stability remains debated [6-8]. Studies on various bacterial species with natural competence have yielded mixed results on the effects of natural transformation on adaptive evolution, suggesting that its role in accelerating bacterial adaptation is not universal and may prove to be beneficial [9, 10], neutral [11], and/or context dependent [10, 12-14]. Despite its contribution to genome evolution, the immediate effects of DNA uptake and recombination on individual cells are not clear. Two non-mutually exclusive hypotheses have been proposed: one suggests that taken-up DNA serves as a source of essential nutrients such as nitrogen, carbon, and nucleotides [6, 8, 14] while the other proposes that it can be used as a template for repair of genomic DNA damage [15, 16]. Experimental evidence supports both hypotheses in different bacterial species, but competence induction may also be a general stress response [17-20].

*Deinococcus radiodurans* is a unique bacterium known for its exceptional ability to survive various DNA-damaging conditions [21]. It employs distinct strategies and DNA repair mechanisms, including RecA-independent and RecA-dependent repair phases [21-33]. *D. radiodurans* cells are naturally transformable, and their competence is maintained throughout the exponential growth phase [34]. DprA, DdrB, and RecO/RecF proteins play important roles in the natural transformation process of *D. radiodurans* [35, 36]. While DprA and RecA is crucial for both plasmid and chromosomal DNA transformation, DdrB and RecO/RecF have distinct roles in this process [35, 37]. Despite the early discovery of the natural transformation phenotype in *D. radiodurans*, there has been limited research on the underlying mechanism and its significance in DNA damage repair and radiation survival. Recent studies have highlighted the role of DprA in protecting and facilitating the integration of incoming DNA into the bacterial chromosome [35, 37]. However, the exact contribution of DprA in DNA repair and its involvement in double-strand break repair mechanisms remains unclear.

In this study, we aimed to elucidate the specific roles of DprA in DNA double-strand break repair and provide compelling evidence demonstrating its importance in balancing the activities of SSA and ESDSA repair pathways.

## Results

### 1. Natural transformation specific genes mutant showed differential response to DNA damaging agents

We investigated the cellular survival of *D. radiodurans* mutants lacking natural transformation specific genes under various DNA damaging conditions. The *comEA* and *pilT* mutants displayed similar cell survival as the wild type when exposed to gamma radiation, UVC radiation, and Mitomycin C (MMC) (Fig. 1, S1, and S2). In contrast, the *ΔendA* mutant exhibited slight sensitivity to gamma radiation (Fig. 1), while MMC treatment leading to a significant reduction in survival (∼1 log cycle drop with MMC at 20 μg/ml for 30 minutes) (fig. S1). UVC treatments did not affect the survival of *ΔcomEA* and *ΔpilT* mutants, but *ΔendA* mutants showed sensitivity (Fig. S2).

**Figure 1:**
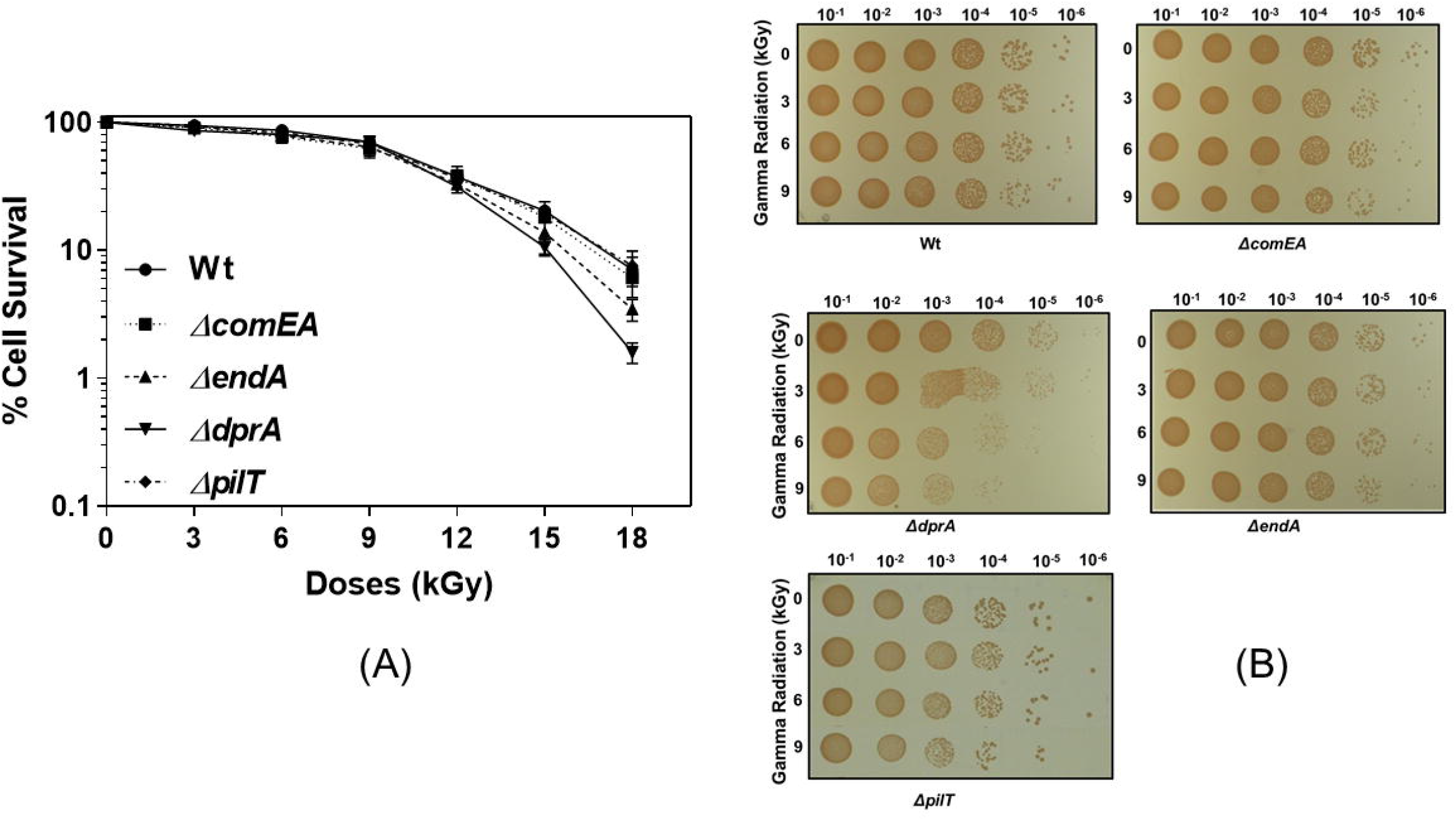
The cell survival of *D. radiodurans* and its mutants after exposure to gamma radiation. (A) Exponential growth phase cells of wild type (-●-), *ΔcomEA* (-■-), *ΔendA* (-▲-), *ΔdprA* (-▼-), and *ΔpilT* (-♦-) mutants were subjected to different doses of gamma radiation, and the percentage of cell survival fraction was plotted as a function of the gamma radiation doses (kGy). The mean ± SEM of three independent experiments were shown. (B) Exponential growth phase cells of wild type and different mutant cells were exposed to gamma radiation doses ranging from 0 to 9kGy. After dilution, aliquots were placed on TGY medium plates and incubated at 32^0^C for 48 hours.

The *ΔdprA* mutant displayed survival rates comparable to the wild type up to 10kGy doses of gamma radiation. However, at 18kGy, the *ΔdprA* cells exhibited nearly a 1-log cycle drop in survival compared to the wild type (Fig. 1). Furthermore, the *ΔdprA* cells demonstrated a 1.3-log cycle drop in survival upon treatment with MMC (20 μg/ml) for 30 minutes (Fig. S1). UVC radiation resulted in a reduction of cell survival by approximately 0.5-log cycle at a dose of 1000 J m^-2^ (Fig. S2). The normal growth patterns of the wild type and different mutants (*ΔcomEA*, *ΔdprA*, and *ΔendA*) were nearly identical, indicating that these gene deletions had no effect on the normal growth of *D. radiodurans* (Fig. S3, A). However, when exposed to gamma radiation (6 & 12kGy), *ΔdprA* and *ΔendA* mutants exhibited some growth defects, possibly due to their increased sensitivity to higher radiation doses (Fig. S3, B & C).

### 2. Analysis of DSB repair in natural transformation (NT) gene mutants

DSB repair of *ΔcomEA*, *ΔdprA*, and *ΔendA* mutants were analyzed by pulse field gel electrophoresis (PFGE). The DSB repair pattern of the *ΔcomEA* mutant was found to be identical to that of the wild type. In contrast, the *ΔendA* mutant exhibited slightly slower DSB repair kinetics compared to the wild type when exposed to 6kGy gamma radiation (Fig. 2A). Interestingly, the *ΔdprA* mutant displayed faster DSB repair kinetics compared to the wild type and within 1 hour of post-Irradiation recovery (PIR) time, approximately eleven distinct bands appeared on the gel after genome digestion with NotI, whereas the corresponding pattern for the wild type appeared on the gel after 2 hours of PIR (Fig. 2B). These findings indicate the involvement of DprA in DSB repair and suggest that its absence results in accelerated repair kinetics. It is noteworthy that both the *ΔdprA* and *ΔendA* mutants exhibited sensitivity to higher doses of gamma radiation, although the level of gamma sensitivity varied. Despite this, the DSB repair kinetics differed between the mutants, with the *ΔendA* mutant showing delayed repair (Fig. 2A) and the *ΔdprA* mutant demonstrating faster repair (Fig. 2B). These observations suggest that the mechanisms of DSB repair induced by gamma radiation may differ in these mutants.

**Figure 2:**
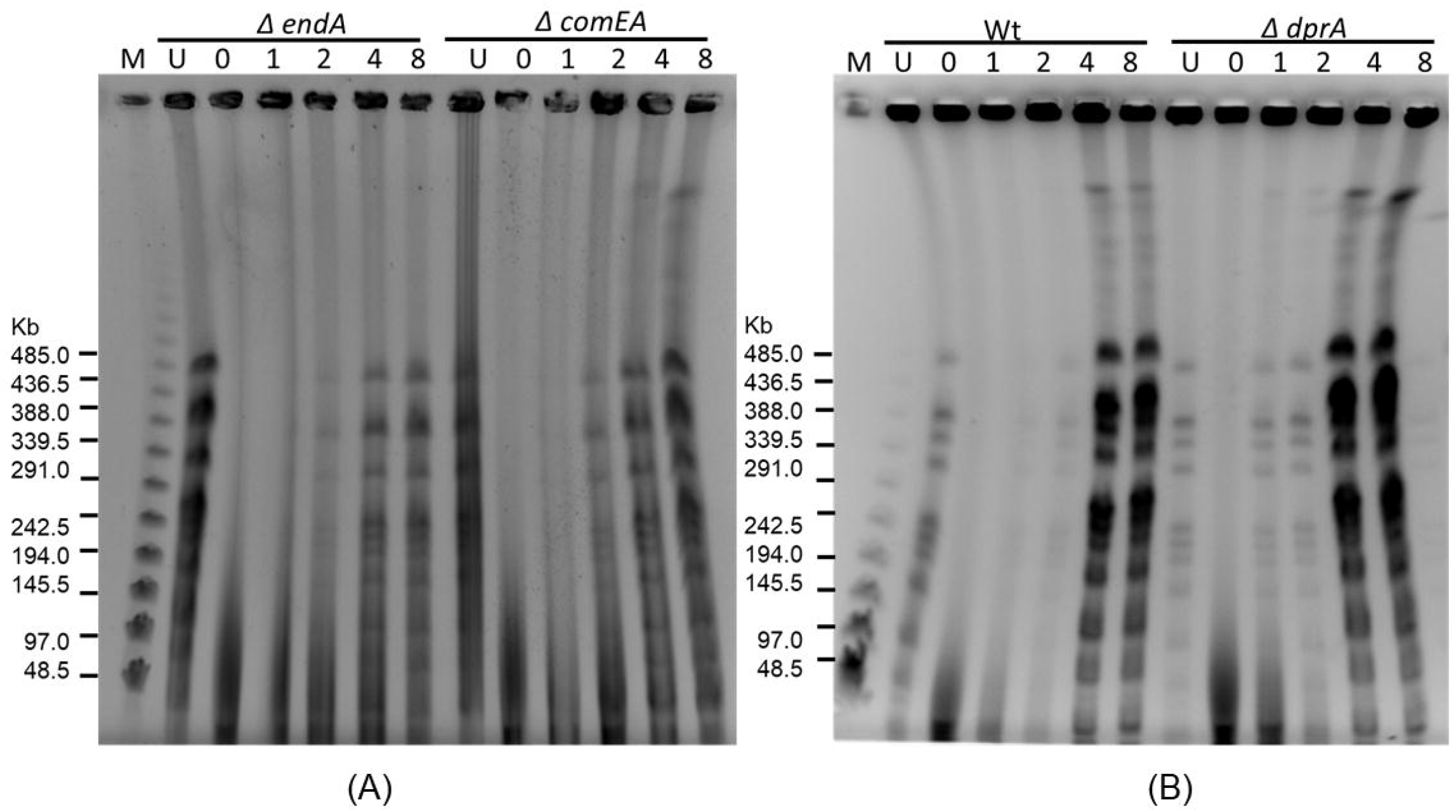
The DNA double-strand breaks (DSBs) repair kinetics of wild-type and natural transformation mutants of *D. radiodurans*. Pulsed-field gel electrophoresis (PFGE) was utilized to evaluate the repair kinetics. (A) The kinetics of DSB repair in *ΔendA* and *ΔcomEA* mutants are shown, while (B) displays the kinetics of DSB repair in wild-type and *ΔdprA* mutant. The NotI-digested DNA from unirradiated cells (U) and irradiated cells at different post-irradiation time points (PIR) after exposure to 6kGy were visualized immediately after irradiation (0) and at specified incubation times (hours) on the gel. Lambda PFG molecular mass standards (lane-M).

Furthermore, the PFGE banding patterns for all studied NT-specific gene mutants were nearly identical to the wild type, indicating the absence of large genomic rearrangements (Fig. 2). Intriguingly, the *ΔdprA* mutant cells also displayed sensitivity when exposed to the DNA-damaging agent MMC (Fig. S1). However, the DSB repair profile of the *ΔdprA* mutant cells showed delayed repair of MMC-induced DSBs compared to wild-type cells, which contrasts with the repair profile observed after exposure to gamma radiation-induced DSBs (Fig. S4). Conversely, the *ΔendA* mutant exhibited delayed DSB repair for both MMC and gamma radiation-induced DSBs (Fig. 2A). These findings suggest that DSBs induced by different agents may follow distinct repair strategies, highlighting the complexity of DSB repair mechanisms in *D. radiodurans*.

### 3. *ΔdprA* mutant show a limited contribution of ESDSA repair

DSBs considered as severe form of DNA damage and can lead to chromosomal abnormalities, genomic instability, and cell death [38]. To counteract these effects, cells have developed several mechanisms for DSB repair, including non-homologous end joining (NHEJ), single-strand annealing (SSA), synthesis-dependent strand annealing (SDSA), break-induced replication (BIR), interstrand SSA, and copy choice. However, none of these mechanisms fully explain the observed DSB repair in *D. radiodurans* [39]. A unique patchwork-type DSB repair mechanism, called ESDSA, has been proposed for *D. radiodurans*. This repair mechanism involves the assembly of DNA fragments in a patchwork pattern to restore the genome [21, 39, 40].

The faster DSB repair kinetics observed in the *ΔdprA* mutant compared to wild-type cells raised questions about the involvement of ESDSA repair in this mutant. To investigate this, we performed ESDSA repair experiment by BrdU labeling followed by UV photolysis of repaired DNA in the *ΔdprA* mutant as describe elsewhere [21, 39, 40]. Surprisingly, the results showed that the *ΔdprA* mutant cells exhibit lesser contribution of ESDSA repair, as evidenced by the absence of DSBs in the repaired genome after reconstitution in BrdU medium and subsequent UV photolysis (Fig. 3A). In contrast, wild-type cells displayed the characteristic patchwork repair pattern associated with ESDSA repair (Fig. 3A, compare lanes 11 and 12). The PFGE pattern of both wild-type and *ΔdprA* mutant cells remained the same in unirradiated, irradiated, and cells recovered for 3 hours PIR or BrdU-labeled cells not exposed to UV photolysis (Fig. 3A, lanes 1-6, and 9-10). Additionally, neither wild-type nor *ΔdprA* mutant cells recovered in BrdU-free medium exhibited DSBs upon UV photolysis (Fig. 3A, lanes 7 and 8). To further investigate, we conducted a UV dose-dependent experiment. Both wild-type and *ΔdprA* mutant cells recovered in BrdU medium were exposed to incremental doses of UV radiation. The results demonstrated that wild-type cells exhibited increasing sensitivity to UV doses (100 to 3000 J m^-2^), with incremental damage and generation of DSBs in the genome. In contrast, the genome of *ΔdprA* mutant cells did not display any UV photolysis sensitivity until exposed to a UV dose of 1000 J m^-2^. However, further increases in UV doses led to the formation of DSBs and genome degradation in the *ΔdprA* mutant cells (Fig. 3B). These findings provide evidence for the limited involvement of ESDSA repair for DSBs repair in *ΔdprA* mutant and suggest a role of *dprA* gene in possible regulation of the choice of DSB repair pathways in *D. radiodurans*.

**Figure 3.**
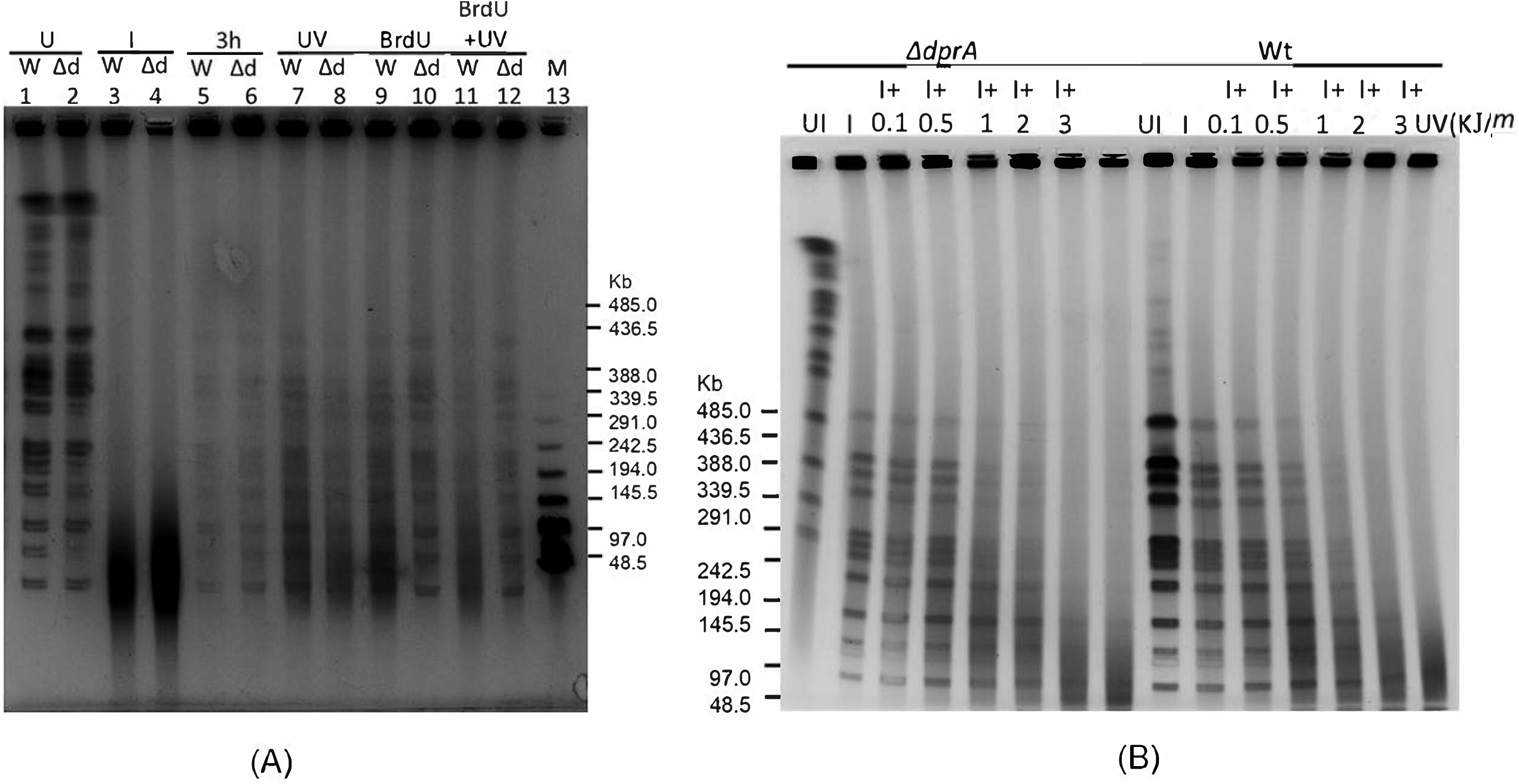
The UV photolysis of the genome of *D. radiodurans* repaired in the presence of BrdU. (A) The NotI restriction pattern of wild-type *D. radiodurans* (W) and *ΔdprA* mutant (Δd) (lanes 1-12) is displayed, along with Lambda PFG molecular mass standards (lane 13). The DNA from cells that were unirradiated (lane 1, 2), irradiated cells (6kGy) immediately after exposure (0 hours PIR) (lane 3, 4), irradiated cells incubated for 3 hours in TGY medium (lane 5, 6), irradiated cells incubated for 3 hours in TGY medium and exposed to 1,000 J m^-2^ of ultraviolet light (UV) (lane 7, 8), and irradiated cells repaired in BrdU-supplemented TGY medium prior (lane 9, 10) or after (lane 11, 12) exposure to 1,000 J m^-2^ of UV are shown. (B) The UV photolysis of cells of wild type and *ΔdprA* recovered in BrdU-supplemented TGY medium post-irradiation is presented. The DNA from unirradiated cells (UI) and 6 kGy gamma irradiated cells (I) repaired in BrdU-supplemented TGY medium for 3 hrs followed by UV exposure (0.1, 0.5, 1, 2, and 3 kJ m^-2^). Lambda PFG molecular mass standards (lane-M)

### 4. Deletion of *dprA* gene in *ΔddrB* mutant improves gamma radiation cell survival and recovery

*D. radiodurans* utilizes SSA and ESDSA type DSB repair mechanisms for repairing its shattered genome. To investigate the role of DprA in regulating the utilization of SSA and ESDSA repair pathways in response to gamma radiation, we conducted studies on radiation survival, DNA synthesis, and DNA degradation in various mutants (*ΔpprA*, *ΔddrA*, and *ΔddrB*) of *D. radiodurans*. Upon exposure to gamma radiation, Deinococcus cells exhibited an upregulation of specific proteins, including SSA repair specific DdrA and DdrB. DdrB has been shown to have multiple functions in cellular processes, such as participating in the repair of genetic instability in repeated sequences through SSA, relieving blocked replication forks, and aiding in early genome repair after exposure to high doses of ionizing radiation [35, 41-46]. DdrA has been demonstrated to assist DdrB in its repair functions. Radiation-induced protein, PprA protein, play a role in DNA repair and cell division. PprA rapidly accumulates at the site of DNA damage, where it forms a complex with other repair proteins to stabilize the broken ends of the DNA and facilitate efficient repair [47, 48]. The observed decrease in ESDSA-mediated repair in the absence of DprA implies that DprA may either directly participate in the ESDSA pathway or indirectly regulate its activation as illustrated in Figure 3.

To investigate the influence of DprA on SSA and ESDSA repair pathways, we deleted the *dprA* gene in *D. radiodurans* strains lacking the *pprA, ddrA*, and *ddrB* genes and examined DSB repair and cell survival in these double mutants. The results showed that the *ΔdprA* and *ΔddrA* single mutants were only sensitive to high doses of gamma radiation (>10 kGy), consistent with expectations. Similarly, the *ΔpprA* and *ΔddrB* mutants exhibited high sensitivity to gamma radiation (Fig. 4). The *ΔddrA ΔddrB* double mutant showed slightly lower survival rates compared to the *ΔddrB* single mutant (Figure 4A and C). However, the deletion of the *dprA* gene in the *ΔddrB* or *ΔddrA ΔddrB* background significantly increased cell survival (∼1.5 log cycle) compared to the *ΔddrB / ΔddrA ΔddrB* mutant alone (Fig. 4A and C). This finding suggests that *dprA* and *ddrB* genes do not exhibit epistasis, indicating that their functions are not mutually dependent. Moreover, this observation also suggests that DprA may interfere with ESDSA repair, which is the sole active DSB repair pathway in the absence of the *ddrB* gene (*ΔddrB* mutant). The growth curve analysis further supported these findings. Wild type and different mutants displayed similar growth patterns when no gamma radiation given (fig. S5, A). However, gamma irradiated *ΔddrB* cells required a considerable amount of time (>1000 min PIR) to recover from the stress induced by gamma radiation, whereas the *ΔddrB ΔdprA* mutant cells initiated growth after 400 min of PIR (Fig. S5 B, C). Besides this, *ΔdprA* cells exhibited a shorter lag phase and resumed growth earlier compared to wild-type cells (Fig. S5 B, C). Together, these data suggest that SSA repair specific *ddrB* gene deletion renders *D. radiodurans* cells sensitive to gamma radiation and the deletion of the *dprA* gene in the *ΔddrB* mutant positively impacts cell survival and recovery from gamma radiation-induced stress possibly by influencing ESDSA repair in *ΔddrB* cells. We also investigated the effect of deleting the *dprA* gene in the *ΔpprA* genetic background and found that it had no impact on gamma radiation cell survival, which remained comparable to that of the individual single mutants (Fig. 4B, C) suggesting epistasis relation of these genes. However, the *ΔpprA ΔddrA* double mutant more sensitive than *ΔpprA* mutant alone for MMC survival and exhibited a DSB repair defect in PFGE gel analysis (Fig. S4).

**Figure 4:**
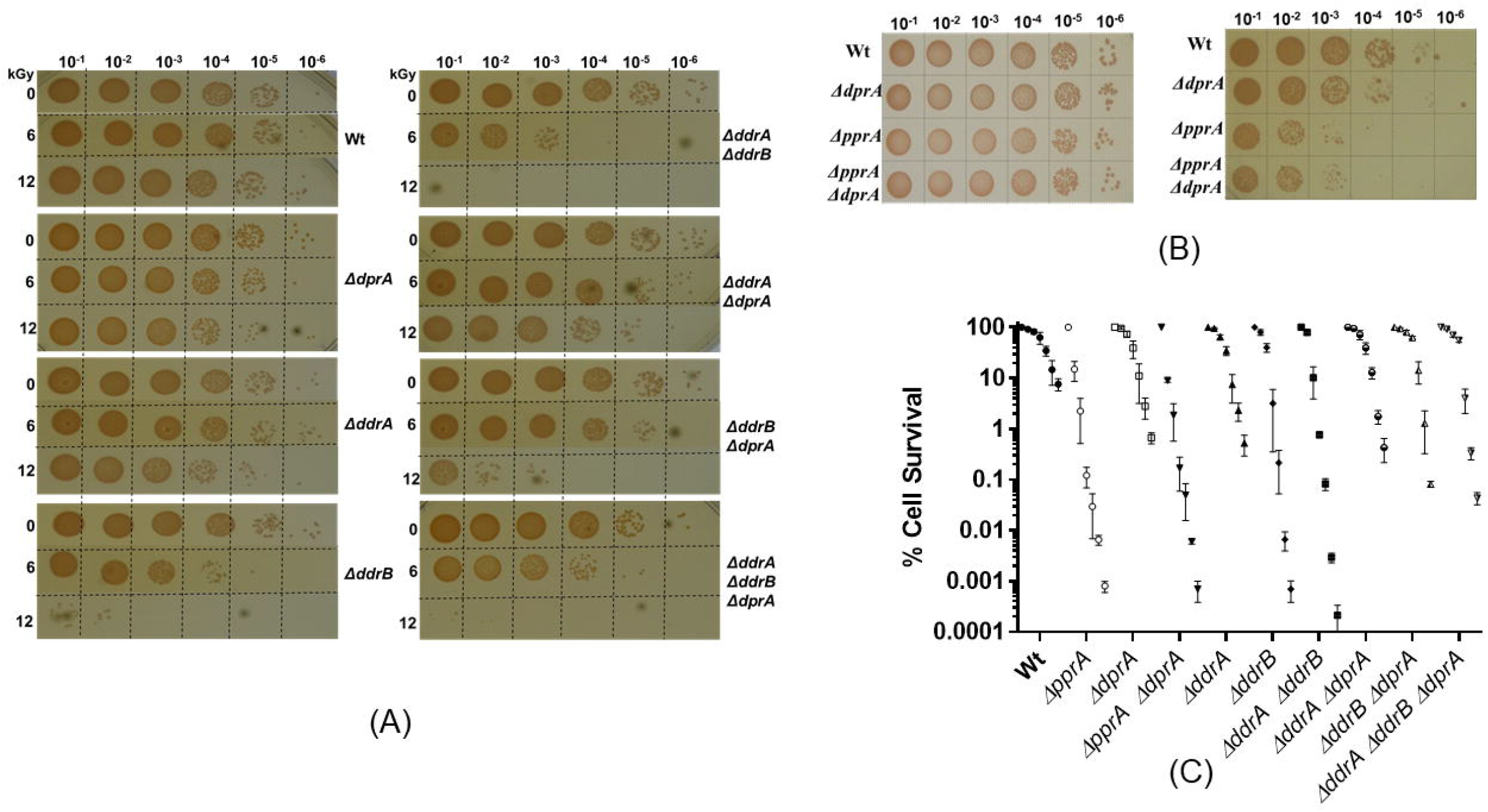
Gamma radiation survival of various mutants of *D. radiodurans*. (A) Exponential growth phase cells of wild type and mutants including *Δdp*rA, *ΔddrA*, *Δddr*B, *ΔddrA ΔddrB*, *ΔddrA ΔdprA*, *ΔddrB ΔdprA*, and *ΔddrA ΔddrB ΔdprA* were exposed to gamma radiation doses ranging from 0 to12kGy. After dilution, aliquots were spotted on TGY medium and incubated at 32°C for 48 hours. (B) Wild type and mutants including *ΔdprA*, *ΔpprA* and *ΔpprA ΔdprA* were exposed to 9kGy gamma radiation doses. After dilution, aliquots were spotted on TGY medium and incubated at 32°C for 48 hours. (C) Exponential growth phase cells of various *D. radiodurans* R1 mutants and wild type were exposed to different doses of gamma radiation, and the percent cell survival fraction was plotted as a function of the radiation doses (0, 3, 6, 9, 12, 15, and 18 kGy). The wild type (-●-) and mutants included *ΔpprA* (-O-), *ΔdprA* (-□□-), *ΔdprA ΔpprA* (-▼-), *Δddr*A (-▲-), *ΔddrB* (-♦-), *ΔddrA ΔddrB* (-■-), *ΔddrA ΔdprA* (- ◒ -), *ΔddrB ΔdprA* (- ◭ -) and *ΔddrA ΔddrB ΔdprA* (- ⧩ -), represented. The mean ± SEM of three independent experiments were shown.

### 5. DprA has inhibitory role on ESDSA Repair Pathway in *D. radiodurans*

In order to better understand the inhibitory role of the DprA protein on the ESDSA repair pathway, we examined the process of DNA end resection involved in DSB repair. [^3^H]-thymidine prelabeled genome analysis revealed minimal DNA degradation in the absence of gamma radiation exposure, with similar levels observed across the wild type and various mutants (within a statistical variation of 10%) (Fig. 5A). However, upon exposure to a 6 kGy dose of gamma radiation, the wild type exhibited the characteristic DNA degradation pattern, initiating promptly after radiation treatment and persisting for 1.5 hours post-irradiation time (PIR) (Fig. 5B). In contrast, the *ΔdprA* mutant displayed a brief DNA degradation phase of 0.5 hours. Notably, the *ΔddrB ΔdprA* double mutant, which exhibited improved gamma survival compared to the *ΔddrB* alone mutant, showed a DNA degradation pattern similar to the wild type but for a reduced duration of approximately 1 hour and with less DNA degradation than the *ΔddrB* alone mutant (Fig. 5B). The *ΔpprA ΔdprA* mutant demonstrated different results compared to the *ΔpprA* alone mutant, as DNA degradation was absent until 2 hours PIR in both mutants, then *ΔpprA* showed DNA degradation but the degradation phase continued until 5 hours PIR in the *ΔpprA ΔdprA* mutant (Fig. 5B).

**Figure 5:**
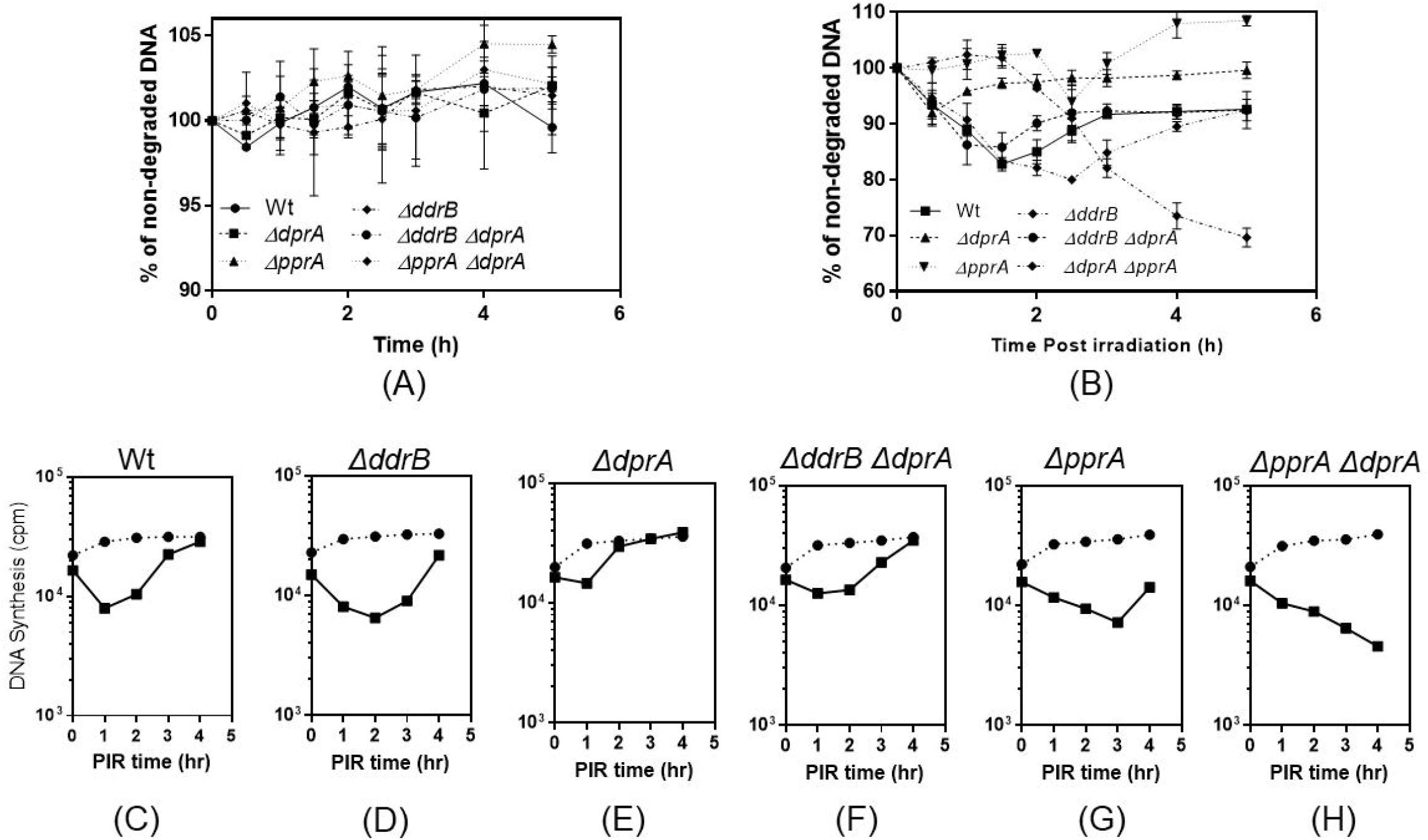
Measurement of DNA degradation and synthesis in wild-type and various mutants. (A) & (B) The DNA degradation was measured in [^3^H]-thymidine prelabeled unirradiated (A) and 6 kGy-irradiated (B); wild type and different mutants at different PIR or respective sham control, and the data are represented as mean ± SEM of two independent experiments. (C to H) The global rate of DNA synthesis was measured by incorporating [3H]-thymidine during a 15-min pulse labeling in 6 kGy-irradiated wild type and various mutants cells, and the results are presented as counts per minute (cpm).

ESDSA repair is characterized by DNA degradation followed by extensive DNA synthesis during repair following irradiation, while SSA repair does not involve significant DNA synthesis [39-41, 49]. In our study, we evaluated DNA synthesis data in wild-type cells and different mutants using a [^3^H]-thymidine pulse for 15 minutes at various PIR time points (Fig. 5C to H). Concurrently, we examined the kinetics of DSB repair using PFGE (Fig. S6). Our results showed that DNA synthesis in wild-type cells occurred after 1.5 hours PIR (Fig. 5C). However, DNA synthesis was considerably delayed (>3 hours) in the *ΔpprA* and *ΔpprA ΔdprA* double mutants (Fig. 5G, H), while in the *ΔddrB* mutant, DNA synthesis commenced after 2 hours PIR (Fig. 5D). Intriguingly, the *ΔdprA* mutant exhibited an early onset of DNA synthesis after a brief period (0.5 hours PIR) (Fig. 5E), whereas the *ΔddrB ΔdprA* mutant initiated DNA synthesis at ∼1.5 hours PIR (Fig. 5F). The delay in DNA synthesis observed in the *ΔpprA* and *ΔddrB* mutants correlated well with their gamma radiation sensitive phenotype and DSB repair kinetics, as evidenced by the NotI pattern on the PFGE gel and consistent with earlier studies (Fig. S6). Furthermore, the early repair of DSBs in the *ΔdprA* mutant corresponded to the onset of early DNA synthesis in this mutant (Fig. 2B & 5E). Additionally, the *ΔddrB ΔdprA* mutant exhibited an earlier pick-up in DNA synthesis compared to the *ΔddrB* mutant, suggesting that the presence of DprA in the *ΔddrB* mutant may have an inhibitory effect on ESDSA repair, as the *ΔddrB* mutant relied on the ESDSA pathway for DSB repair due to the absence of the functional SSA pathway. To provide further support for these findings, cell survival and growth analysis data of both the wild type and *ΔdprA* mutant strains overexpressing DprA from the pRAD plasmid done (Fig. S7). Notably, a dominant negative effect on cell survival and growth curve was observed in wild type cells overexpressing DprA, particularly at gamma radiation doses tested (6 &12 kGy). A similar effect was also observed in the *ΔdprA* mutant strain overexpressing DprA, albeit with less severity due to the absence of chromosomal doses of DprA (Fig. S7). In conclusion, data suggest that DprA protein plays an inhibitory role in the ESDSA repair pathway of *D. radiodurans* and overexpression of DprA negatively affects cell survival and growth.

### 6. DprA may sequester RecA focus *in vivo*

RecA Protein Plays a Crucial Role in the ESDSA Repair Pathway in *D. radiodurans*. RecA loading onto ssDNA facilitated by RecFOR complex as RecBC complex is absent in *D. radiodurans* [50]. DprA is proposed to bind to ssDNA and protect it from degradation by nucleases during natural transformation [51] and facilitate RecA loading during DNA recombination between different bacterial species [36]. The cost and benefit of transforming DNA during natural transformation are subjective and context-specific. However, under genotoxic stress, transforming DNA may introduce additional DNA strand breaks, potentially negating the beneficial effects such as the use of transforming DNA as a nutritional source [52]. Therefore, the observed inhibitory effect of DprA in ESDSA repair could be explained by the hypothesis that during early ESDSA repair, the RecFOR complex facilitates RecA loading onto ssDNA. However, due to its recombination mediator-like activities, DprA may compete with RecFOR to load RecA protein onto ssDNA generated through radiation-induced ssDNA ends or on transforming DNA, thereby reducing the availability of ssDNA-bound RecA foci capable of catalyzing ESDSA repair.

To test this hypothesis, we evaluated the percentage frequency of RecA^RFP^ foci formation in wild-type, *ΔddrB*, and *ΔdprA* mutants. The data showed that RecA foci formation increased immediately after irradiation but gradually returned to the unirradiated level by 4 hours PIR. In *ΔddrB* mutant RecA foci formation elevated at 4 hours PIR. However, this dynamic pattern was not observed in the *ΔdprA* mutants until 4 hours PIR (Fig. 6 & S8). Furthermore, in wild-type cells, one-way ANOVA analysis of the 0-hour PIR samples from the wild type, *ΔdprA*, and *ΔddrB* showed a significant difference in RecA-foci formation (*P* = 0.006) (Fig. 6). These findings suggest that DprA may actively facilitate the formation of the RecA nucleoprotein complex with ssDNA, potentially due to its recombination mediator-like activities. However, it remains unclear whether RecA foci formed with DprA-loaded ssDNA are proficient in recombination repair or defective. Nevertheless, the microscopy data, in conjunction with the findings on cell survival and DSB repair kinetics presented earlier, strongly suggest that DprA exerts its function by diminishing the effective availability of RecA-ssDNA complex molecules for the RecFOR-mediated ESDSA recombination repair pathway. This reduction ultimately leads to a decelerated progression of the ESDSA repair process.

**Figure 6:**
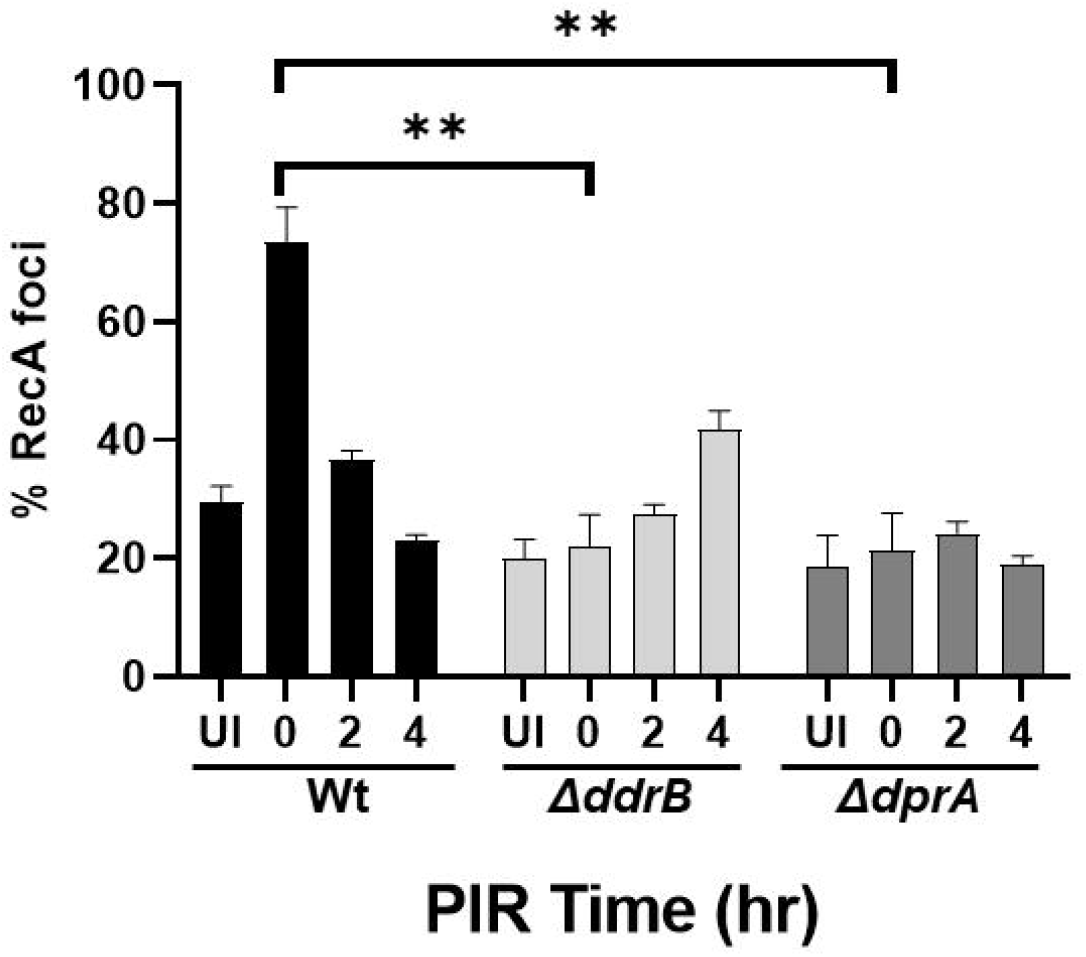
Frequency of RecA foci formation in wild type and different mutants. Fluorescence microscopy images of *D. radiodurans* cells expressing DrRecA^RFP^ in logarithmic growing wild-type and mutant strains (*ΔdprA* and *ΔddrB*). The expression of DrRecA was visualized in the RFP channel (561nm) and under bright field (DIC). Error bars show the S.E.M, and asterisks indicate statistically significant differences when 0hr PIR compared to unirradiated cells (One-way ANOVA, *P* = 0.006).

## Discussion

*D. radiodurans*, known for its exceptional resistance to extreme conditions, possesses distinct repair mechanisms that efficiently mend DNA fragments in to functional genome. The bacterium employs two primary pathways for repairing DSBs; the RecA-dependent extended synthesis-dependent strand annealing (ESDSA) pathway and the RecA-independent single-strand annealing (SSA) pathway. The SSA pathway plays a crucial role in the early repair of fragmented genomes, while the ESDSA pathway involves DNA degradation and extensive synthesis to accomplish patch-type repair [21, 39-42, 49]. SSA repair reduces the number of small DNA fragments by transforming them into larger fragments suitable for ESDSA-mediated homologous recombination repair [41, 49]. The SSA pathway in *D. radiodurans* relies on *DdrB and DdrA* proteins. DdrB aids in DNA annealing during SSA repair, while DdrA helps to limit DNA degradation by nucleases [41, 42, 46]. The presented data in this study emphasizes the regulatory role of the NT-specific protein DprA in governing the choice between DSB repair pathways (SSA and ESDSA) in *D. radiodurans*. Cell survival experiments revealed distinct susceptibility of *ΔdprA* and *ΔendA* mutant strains to gamma radiation, UV radiation, and MMC, indicating the vital functions of these genes in cell survival (Fig. 1, S1, and S2). The differential responses of the *ΔendA* and *ΔdprA* mutants to MMC and gamma radiation may be attributed to distinct mechanisms of DSB repair for each agent, as supported by previous studies [33, 53].

Interestingly, data presented in Fig. 2 & 3 reveals that in wild-type cells, both the SSA and ESDSA pathways contribute to the repair of severely damaged DNA. However, in the absence of DprA, the SSA pathway becomes the primary mechanism for repairing DSBs, resulting in diminished efficiency in error-free patchwork ESDSA repair (Fig. 3). During DSB repair, the resected ssDNA can serve as a template for either SSA or ESDSA, depending on factors such as the length of ssDNA tails or the presence of DNA damage in the resected region. The selection of one pathway over the other is influenced by various factors, leading to an interdependent relationship between the two pathways. In *D. radiodurans*, maintaining a balance between the SSA and ESDSA repair pathways is critical for survival during recovery from genotoxic stress [21]. SSA repair plays a pivotal role in initiating ESDSA repair by providing a suitable substrate for the formation of RecA nucleoprotein filaments [41]. However, key repair proteins from both pathways may compete for repair, favoring SSA repair in the early stages of DSB repair [41, 42]. DdrB promotes rapid annealing of complementary strands using short homology, providing a suitable DNA substrate for ESDSA repair [45]. RecA, a crucial protein for ESDSA repair, efficiently searches for homology and facilitates error-free repair [40, 54]. In heavily irradiated *D. radiodurans* cells, small DNA fragments may not serve as optimal substrates for RecA-catalyzed ESDSA repair. Hence, prioritizing the RecA-independent phase becomes necessary for efficient DSB repair.

The diminished role of ESDSA repair in the *ΔdprA* mutant indicates the apparent role of DprA in regulating the choice between SSA and ESDSA repair pathways in *D. radiodurans* exposed to gamma radiation (Fig. 2 & 3). This observation is further supported by cell biology data showing improved survival of the *ΔddrB ΔdprA* double mutant and *ΔddrA ΔddrB ΔdprA* triple mutant compared to the *ΔddrB* and *ΔddrA ΔddrB* mutants (Fig. 4C). Based on our findings, the decrease in ESDSA repair observed in the *ΔdprA* mutant may be attributed to a concurrent increase in SSA repair. Consequently, we anticipate that the deletion of the *dprA* gene in the *ΔddrB* or *ΔddrA ΔddrB* mutant (which lacks SSA repair) would enhance ESDSA repair, resulting in improved cell survival these cells (Fig. 4). This notion is further supported by the relatively faster repair of DSBs in *ΔddrB ΔdprA* mutants compared to *ΔddrB* alone (Fig. S6) and the rapid DNA synthesis kinetics with a short DNA degradation phase observed in the *ΔdprA* and *ΔdprA ΔddrB* mutants (Fig. 5). These observations were further supported by dominant negative effect of DprA overexpression in wild type cells (Fig. S7). Collectively, these findings suggest that DprA may inhibit RecA-mediated ESDSA recombination repair, and its deletion in the *ΔddrB* mutant could reverse this hindrance, resulting in improved survival. The deletion of the *dprA* gene in the *ΔpprA* genetic background had no effect on cell survival, indicating that DprA is less likely affect PprA functions during DSB repair in *D. radiodurans* [47, 48].

The right choice of DSB repair pathway is crucial to preserve genomic stability. For eukaryotes, cell growth phase and availability of undamaged homologous DNA for repair are crucial parameter for the choice of DSB repair pathway. BLM helicase, along with 53BP1 and RIF1, have been shown to key players in the initial stages of DSB repair pathway determination in mammalian cells [55]. In bacteria, RecA proteins act as molecular search engines during recombination repair, facilitating DNA strand exchange between homologous DNA strands for error-free repair [56, 57]. RecA activity is regulated by various RecA mediator proteins (RMPs) that interact with it, aiding in its loading onto single-stranded DNA (ssDNA) or its removal from it. These RMPs include RecF, RecO, RecR, DprA, PprA, RecU, and RecX. The RecFOR and DprA assists in assembling RecA onto ssDNA, while PprA, RecU, and RecX negatively regulate RecA activity to prevent hyper-recombination [3, 36, 54, 58-64]. The formation of RecA foci on ssDNA is significantly reduced in the *ΔdprA* mutant compared to the wild type, and there are no dynamic changes in RecA foci formation during post-irradiation recovery (PIR) (Fig. 6). These findings suggest that DprA, through its RMP-like activity, may able to effectively competes with repair-specific RMPs like RecFOR for the loading of RecA onto SSB-coated ssDNA ends, thereby slowing down the ESDSA repair process. This indirectly enhances the preference for SSA repair. The DprA led additional DSB generation during NT of *Acinetobacter baylyi* post-UV stress recovery suggested [52]. For *D. radiodurans*, the level of RecA increases immediately after cells receive acute doses of gamma radiation [65]. However, RecA role in ESDSA and homologous recombination (HR)-mediated repair begins at a somewhat later time during post-irradiation recovery (∼1.5 hours) [39, 40]. Therefore, it is crucial to control RecA function until SSA repair is active in the early hours of post-irradiation recovery, and the role of DprA in interfering with RecA recombination activity appears to be an effective and attractive mechanism for regulating RecA activity and efficiently utilizing various DSB repair pathways. Based on the findings of this study, a proposed scheme for efficient DSB repair in *D. radiodurans* involves DprA-mediated regulation of SSA and ESDSA repair (Fig. 7). It is suggested that in the *ΔdprA* mutant, the role of ESDSA in DSB repair is reduced, while simultaneously, SSA repair takes on a more significant role in the repair process. These findings not only shed light on DSB repair mechanisms but also indicate that the function of DprA extends beyond natural transformation in bacteria. This expanded role of DprA may partly explain why the *dprA* gene is widespread even in bacteria that lack natural transformation capabilities. Therefore, the involvement of DprA in influencing the early steps of DSB repair pathway choice in *D. radiodurans* is suggested.

**Figure 7:**
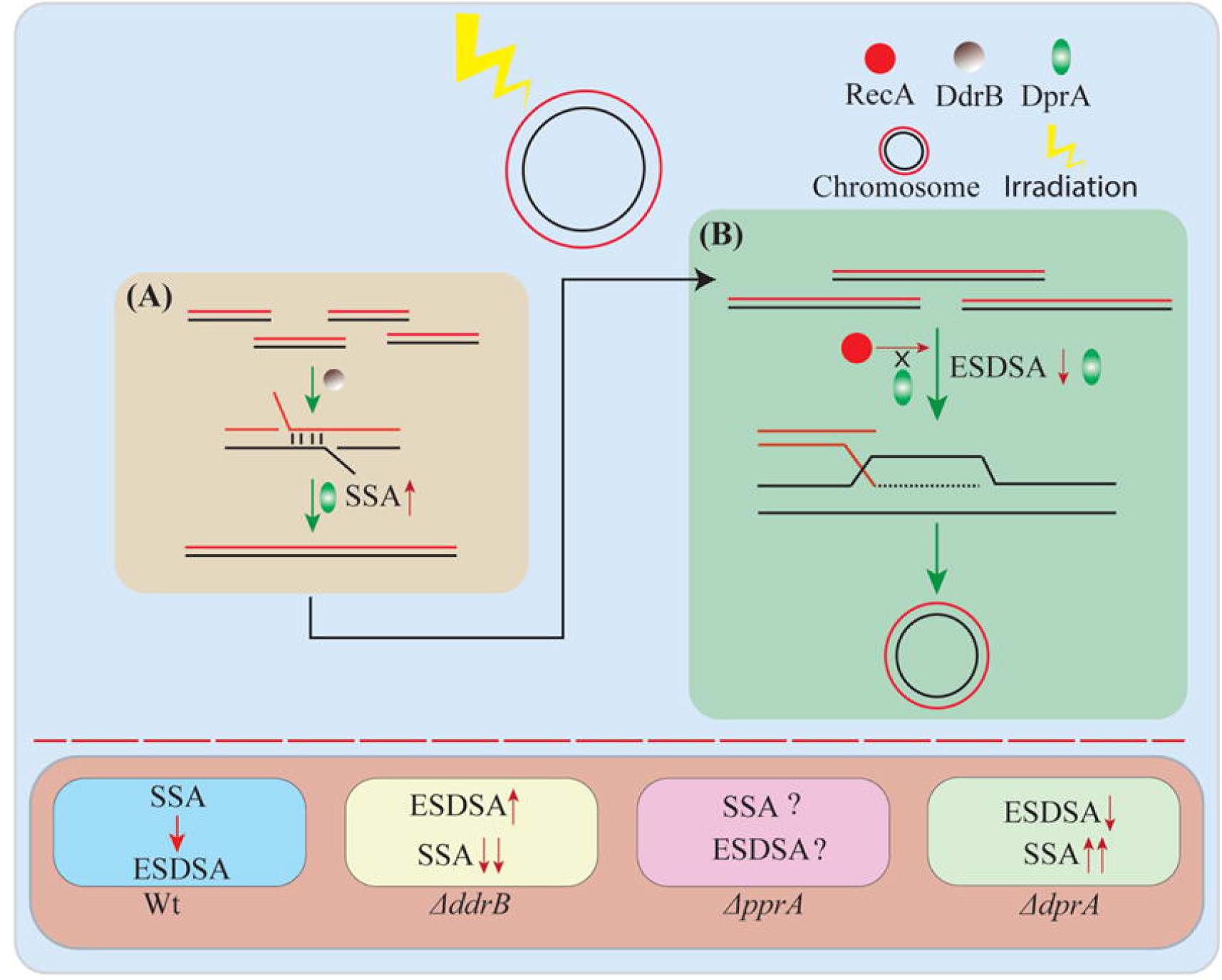
Model explaining the DprA role in DSB repair of *D. radiodurans*. The DSB repair process in *D. radiodurans* proceeds through a RecA-independent phase (A), which is supported by the SSA repair pathway, with crucial involvement of the DdrB and DdrA proteins. During this RecA-independent phase, the DNA fragments are reduced in size to 25-30 kb fragments. These smaller DNA fragments are then joined together with the assistance of ESDSA and HR (homologous recombination) repair mechanisms, resulting in the complete repair of the genome (B). In the lower panel, the relative status of SSA and ESDSA repair is depicted. In the wild-type *D. radiodurans*, SSA repair takes place first, followed by ESDSA repair. However, in the *ΔddrB* mutant, the contribution of SSA repair is diminished, and repair primarily occurs through the ESDSA repair pathway. Conversely, in the *ΔdprA* mutant, the data from the present study suggests that ESDSA role is downplayed, and concurrently SSA repair contributes more significantly to the DSB repair process. As for the *ΔpprA* mutant, the precise share of SSA and ESDSA repair in DSB repair remains unclear.

## Acknowledgements

The authors would like to express their gratitude to Prof. Fabrice Confalonieri and Prof. Pascale Servant from Université Paris-Saclay, CEA, CNRS, Institute for Integrative Biology of the Cell, France, for kindly providing various *D. radiodurans* mutants strain (*ΔddrA*, *ΔddrB*, and *ΔddrA ΔddrB*). Authors further would also like to extend their appreciation to Prof. Issay Narumi from Toyo University, Japan, for sharing the *ΔpprA* mutant. The authors are also thankful to Shri Ranjit Sharma and Shri Lokesh Kumar from BARC for their valuable assistance in scintillation counting.

## Experimental Procedure

### Bacterial strains, plasmids, chemicals, and growth medium

Bacterial strains were employed, including wild type *D. radiodurans* R1 sourced from the ATCC (ATCC 13939), a *ΔpprA* mutant obtained from Prof. I. Narumi, Japan, and *ΔddrA*, *ΔddrB*, and *ΔddrA ΔddrB* double mutants obtained from Prof. Pascale Servant, France. The construction of *ΔdprA*, *ΔendA*, *ΔcomEA*, and *ΔpilT* mutants, as well as the deletion of the *dprA* gene in different genetic backgrounds, was carried out in the laboratory using previously described methods. To isolate the thymine-requiring (thy ^-^) derivatives of the wild type and various mutant strains, they were grown on a solid minimal medium containing thymine (50 μg/ml) and trimethoprim (100 μg/ml) as described elsewhere [39]. All strains were grown in TGY medium (1% Bacto-tryptone, 0.1% glucose, 0.5% yeast extract) supplemented with appropriate antibiotics, as previously described [25]. The media were supplemented with a variety of antibiotics as and when required, including Kanamycin (8 μg/ml), Spectinomycin (75 μg/ml), and Chloramphenicol (3 μg/ml) for *D. radiodurans*. Ampicillin (100 mg/ml) for *E. coli*. The shuttle vector *p11559* was maintained in the presence of Spectinomycin for both *D. radiodurans* (75 μg/ml) and *E. coli* (150 μg/ml), while the shuttle expression vector *pRADgro* and its derivatives were maintained in *E. coli* strain HB101, as previously described [66]. Standard molecular biology techniques were utilized, as outlined in manual (Molecular Cloning: A Laboratory Manual. 4th Edition, Vol. II, Cold Spring Harbor Laboratory Press, New York. Green et al., 2012). Molecular biology grade chemicals, enzymes, and salts were obtained from various sources, including Sigma Chemicals Company, USA; Roche Biochemicals, New England Biolabs (USA); and Merck India Pvt. Ltd., India. Radiolabeled nucleotides were obtained from the Board of Radiation and Isotope Technology (BRIT), Department of Atomic Energy, India. Please refer to Table S1 for a complete list of the bacterial strains, plasmids, and PCR primers employed in this study.

### Cell survival studies

The experimental treatments for *D. radiodurans* cells survival studies included exposure to different doses of UV and gamma radiation as previously described [67], as well as treatment with MMC (20μg/ml) following the protocol described elsewhere [25]. Briefly, bacterial cultures grown in TGY medium at 32°C were washed and suspended in sterile phosphate buffered saline (PBS) and exposed to varying doses of gamma radiation at a dose rate of 5.86kGy per hour (GC-5000, ^60^CO, Board of Radiation and Isotopes Technology, DAE, India). For UVC treatment, different dilutions of cells were plated and exposed to various doses of UV (254 nm) radiation. The treated cells were then plated on TGY agar plates supplemented with appropriate antibiotics, if necessary, and colony-forming units were counted after a 48-hour incubation period at 32°C.

### Measurement of DNA repair kinetics using pulsed-field gel electrophoresis

Irradiated cultures were diluted in TGY to an OD_600_=0.2 and incubated at 32°C. At indicated intervals, 5-ml samples were taken to prepare DNA plugs as described by Mattimore and Battista [68]. The DNA contained in the plugs was digested with 60 units of NotI restriction enzyme (NEB) for 16 hours at 37°C. After digestion, the plugs were subjected to pulsed-field gel electrophoresis in 0.5xTBE using a CHEF-DR® III electrophoresis system (Bio-Rad) at 6 V/cm2 for 20 h at 14°C, with a linear pulse ramp of 50-90 s and a switching angle of 120°.

### Rate of DNA degradation and DNA synthesis measured by ^3^H-thymidine labelling

DNA degradation was estimated as mentioned elsewhere [21, 39, 40]. In brief, pre-label *D. radiodurans* cultures, 10 mCi/ml ^3^H-thymidine (BRIT, DAE, India, specific activity 18 Ci/mmol) was added and the cultures were grown for 18 hours. To eliminate the radioactive thymidine present in the intracellular pools, the cultures were incubated for an additional hour in fresh TGY. Both irradiated and unirradiated cultures were diluted in TGY to an OD_600_ of 0.2 and agitated at 32ºC. Samples of 50 μl were withdrawn at different time points and filtered using Whatman GF/C filters. The filters were then dried and washed twice with 10% TCA, once with 5% TCA, and briefly with 96% ethanol. The nondegraded DNA content was determined by scintillation counting of the dried filters using TRI-CARB 4910TR 110 V Liquid Scintillation Counter, Perkin Elmer.

For the DNA synthesis studies, unirradiated and irradiated exponentially growing cultures were incubated and 0.5-ml samples taken and mixed with a 0.1 ml pre-warmed TGY medium containing 10 μCi ^3^H-thymidine (BRIT, DAE, India, specific activity 18 Ci/mmol). Radioactive pulses were terminated after 15 min by addition of 2 ml ice-cold 10% TCA. Samples were kept on ice for at least 1 hour, and then collected by suction onto Whatman GF/C filters followed by washing with 5% TCA and 96% ethanol. Filters were dried overnight at room temperature, and placed in 5 ml scintillation liquid. The precipitated counts were measured in a liquid scintillation counter.

### BrdU incorporation and UV-induced photolysis of BrdU-substituted DNA

The *D. radiodurans* and its various mutants thy^**-**^ culture were subjected to 6 kGy gamma radiation as described above. The irradiated culture was then diluted to an OD_600_ of 0.2 and grown in TGY supplemented with 5-BrdU for 3 hours. After collecting the cells by centrifugation, they were resuspended in phosphate buffer and starved in the buffer for one hour at 32°C. The cell suspension was then cooled on ice and exposed to various doses of 254-nm UV light in a thin layer. Both UV-irradiated and unirradiated cells were embedded in agarose plugs for subsequent analysis of DNA using PFGE (as described above).

### RecA^RFP^ foci evaluation by Confocal Microscopy

Confocal microscopy was performed on an Olympus FV3000 confocal microscope with a 100x, 1.45 NA oil-immersion apochromatic objective lens. The laser beams were focused on the back focal plane, and the FluoviewTM software was used to control the intensity and time sequence of laser illumination. Fluorescence emission was collected through a DM-405/488 dichroic mirror and single-band emission filters (561 nm). For fixed-cell imaging, bacterial cells in TYG broth were grown to the mid-log phase. They were then fixed with 4% paraformaldehyde on ice for 10 minutes, followed by two washes with PBS. The cells were mounted on glass slides with a 1% agarose bed and observed under the microscope. Image analysis was performed using the cellSens software, analyzing 400-500 cells from at least two separate microscopic fields captured in two independent experiments. The data obtained were analyzed by One-way ANOVA in GraphPad Prism.

### Statistical Methods

The figures display the average values of at least three separate experiments with the standard error of the mean (SEM). For Figure 6, a one-way ANOVA was conducted to determine whether the difference is statistically significant between unirradiated (UI) and 0 hr irradiated samples (*P* =0.006).

