## Supplemental information for "Natural transformation specific DprA coordinate DNA double strand break repair pathways in heavily irradiated *D. radiodurans*"

**Supplementary Data file:**

**Supplementary Data:**

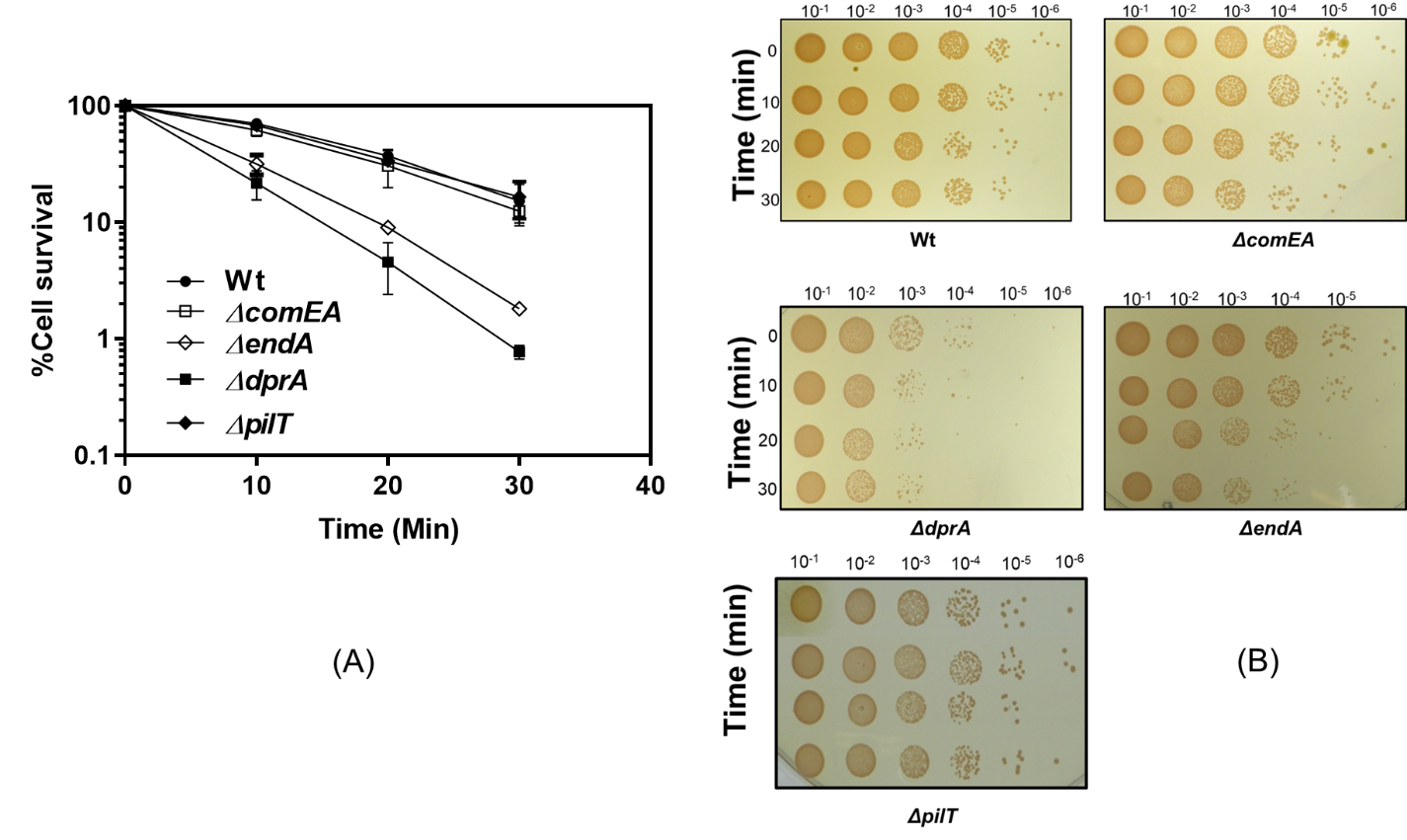

**Figure S1: The cell survival of *D. radiodurans* wild type and its NT mutants after exposure to mitomycin C (MMC).** Exponential growth phase cells of wild type (-●-), *ΔcomEA* (-󠄀󠄀-), *ΔendA* (-◊-), *ΔdprA* (-■-), and *ΔpilT* (-♦-) mutants were mutants were treated with concentrations of 20µg/ml mitomycin C for varying time periods (0, 10, 20, and 30 min), and appropriate dilutions were plated on TGY plates and CFU obtained. The percentage of cell survival fraction was plotted against the treatment time period (min) (A). TGY plates were spotted with appropriate dilutions and cell grown plates were photographed after incubated at 32^0^C for 48 hours (B). The mean ± SEM of two independent experiments were shown.

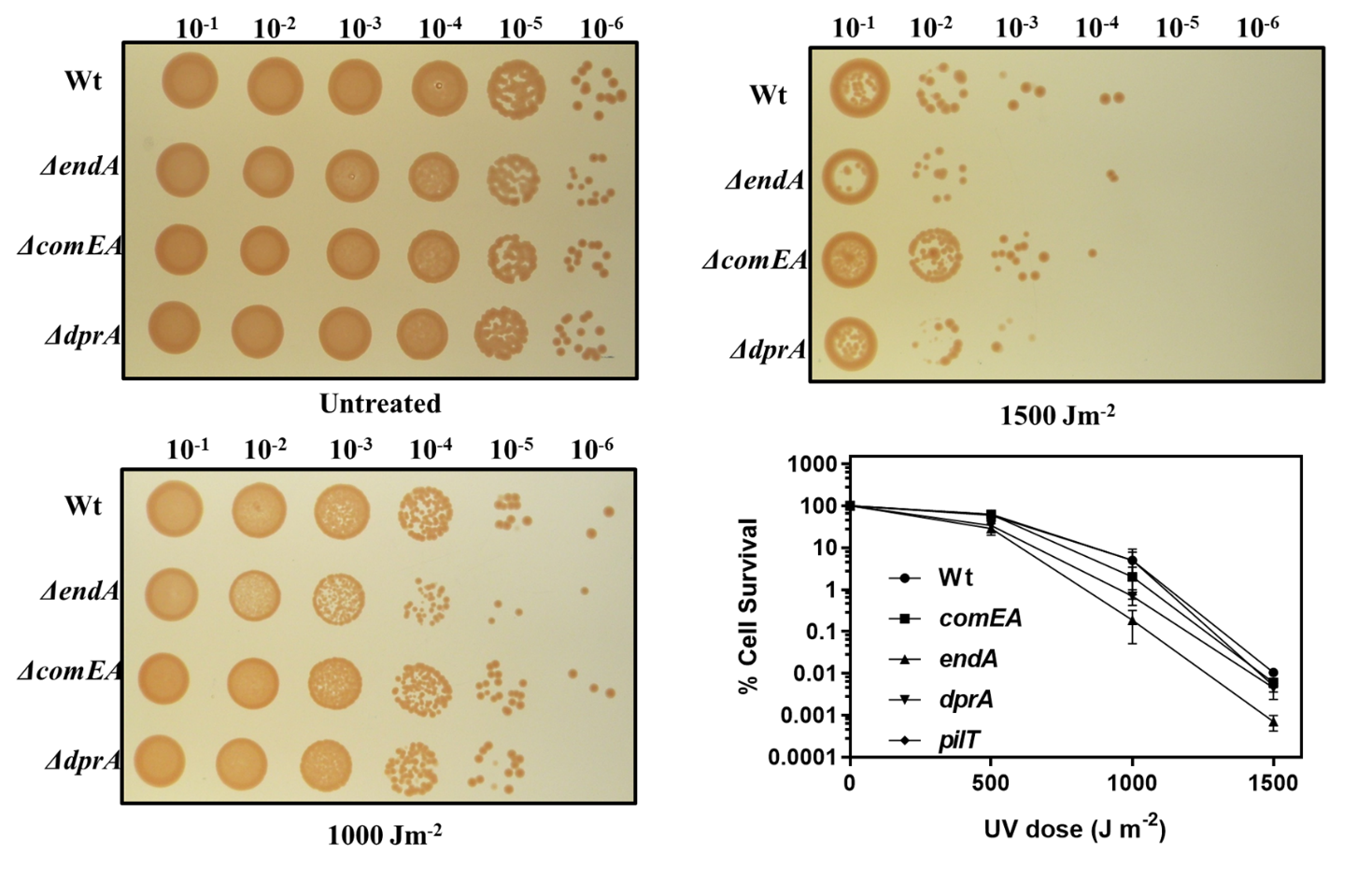

**Figure S2: The cell survival of *D. radiodurans* wild type and its NT mutants after exposure to UV (254nm).** Exponential growth phase cells of wild type (-●-), *ΔcomEA* (-■-), *ΔendA* (-▲-), *ΔdprA* (-▼-), and *ΔpilT* (-♦-) mutants were mutants were treated with different UV doses and after dilution, aliquots were spotted on TGY plates and incubated at 32^0^C for 48 hours. The percentage of cell survival fraction was plotted against the UV dose (J m^-2^). The mean ± SEM of two independent experiments were shown.

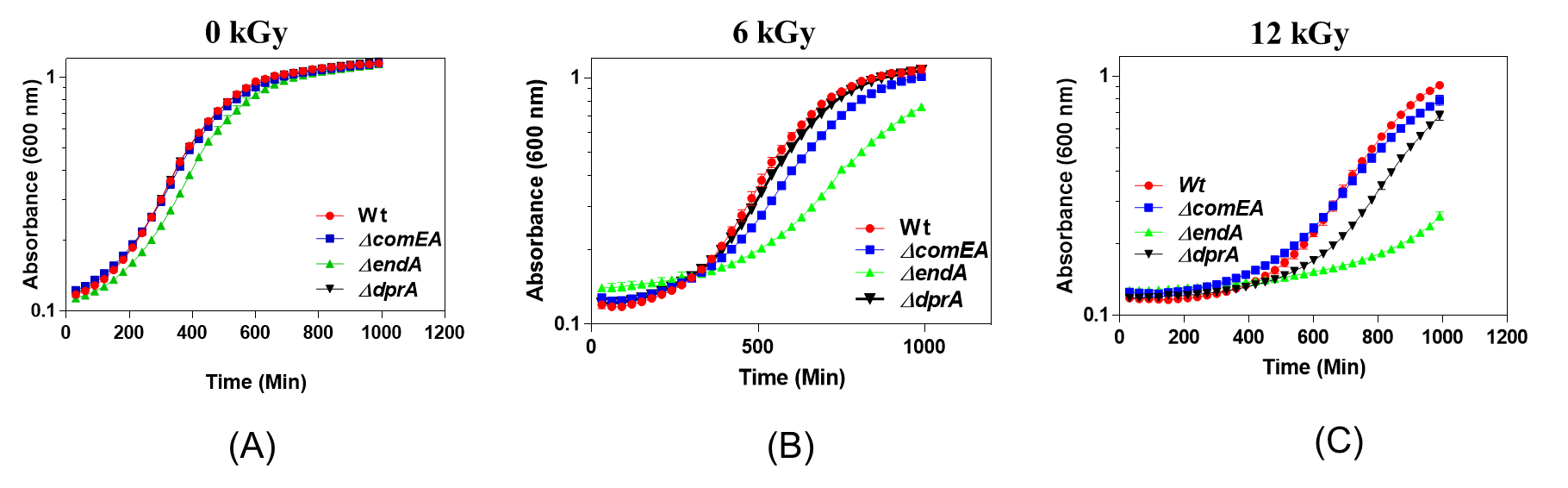

**Figure S3 Cell viability and growth curve of wild type and its NT mutants after exposure to gamma radiation.** Optical density at 600nm was measured continuously using a microtitre-based density reader, and the growth medium (TGY) served as a blank for the entire incubation period. The obtained data was normalized with the blank optical density. Panel (A) shows the growth of normal, untreated cells, panel (B) displays cells treated with 6 kGy of gamma radiation, and panel (C) illustrates cells treated with a dose of 12 kGy of gamma radiation.

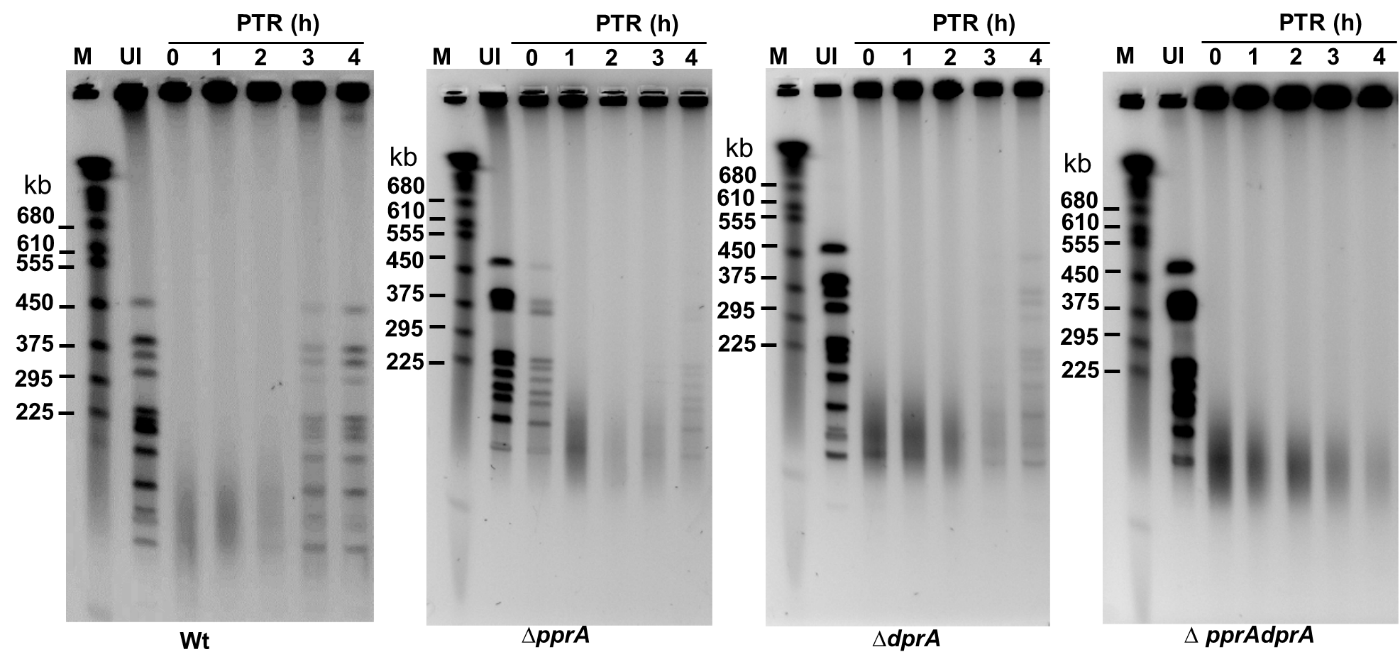

**Figure S4: The repair kinetics of DNA double-strand breaks (DSBs) in *D. radiodurans* wild-type and mutant strains upon treatment with MMC.** The kinetics of DSB repair are shown for four strains, namely wild type (Wt), *ΔpprA*, *ΔdprA*, and *ΔpprA ΔdprA* mutants. The evaluation of the repair kinetics was performed using pulsed-field gel electrophoresis (PFGE). The NotI-digested DNA from untreated cells (UI) and cells treated with 20µg/ml mitomycin C (MMC) for 30 min were allowed to recover in TGY medium for 4 hours post-treatment (PTR). Cell samples collected at different time points were analyzed on PFGE to assess the kinetics of DSB repair. *S. cerevisiae* molecular mass standards (lane-M).

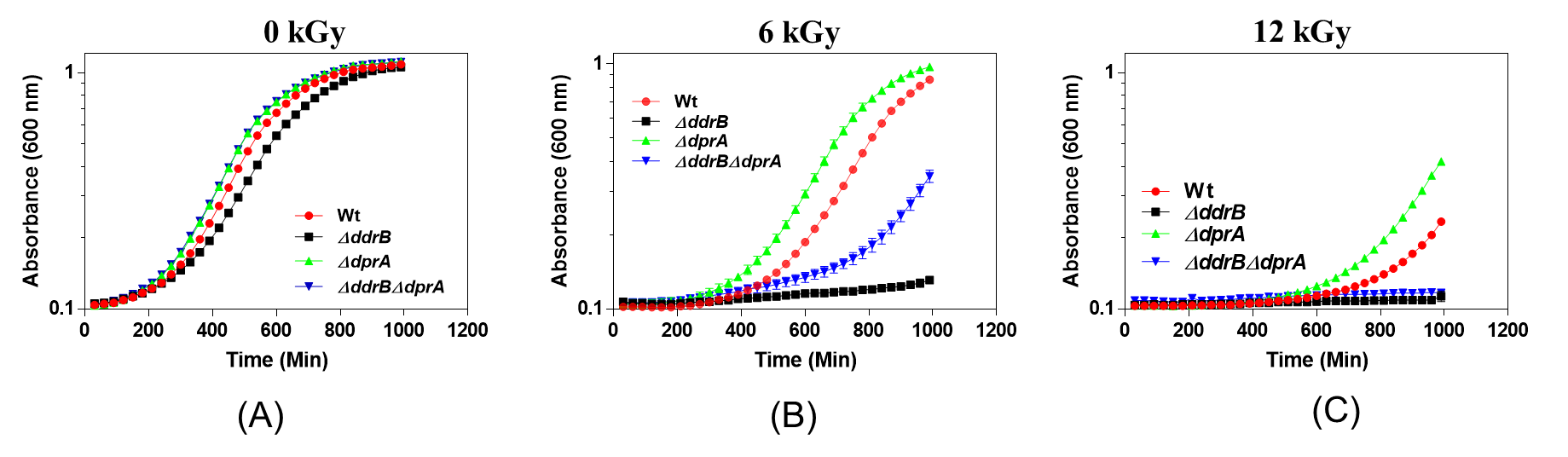

**Figure S5 Cell viability and growth curve of wild type and its mutants after exposure to gamma radiation.** Optical density at 600 nm was continuously measured using a microtitre-based density reader. Panel (A) shows the growth of normal, untreated cells, panel (B) displays cells treated with a dose of 6 kGy of gamma radiation, and panel (C) illustrates cells treated with a dose of 12 kGy of gamma radiation. The data suggests that the deletion of the *dprA* gene in the *ΔddrB* mutant of *D. radiodurans* results in improved cell survival and growth after gamma radiation treatment.

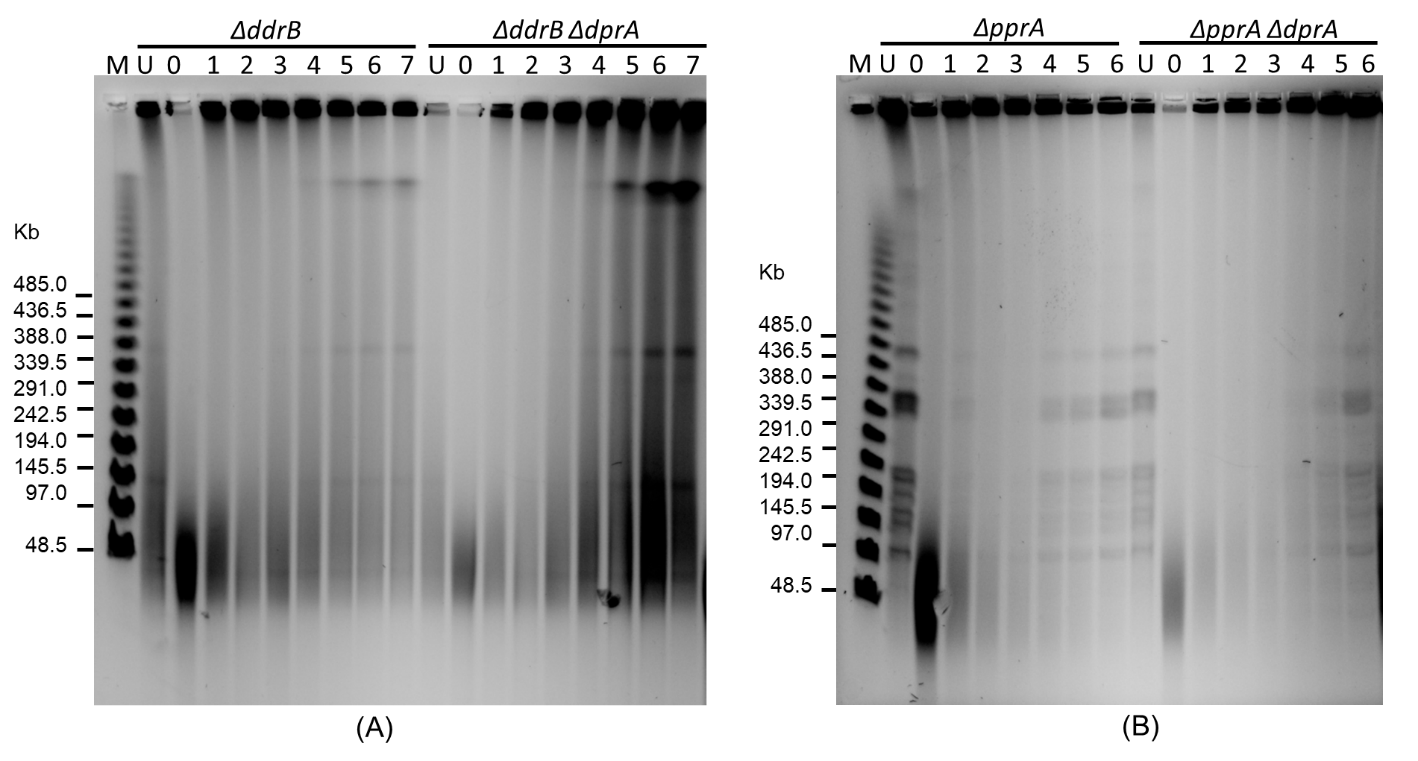

**Figure S6: The DNA double-strand breaks (DSBs) repair kinetics of *D. radiodurans* wild-type and its mutants.**

PFGE was employed to assess the repair kinetics of DSBs. Panel (A) presents the repair kinetics of DSBs in *ΔddrB* and *ΔddrB ΔdprA* mutants, while panel (B) displays the repair kinetics of DSBs in *ΔpprA* and *ΔpprA ΔdprA* mutants. The NotI-digested DNA samples obtained from unirradiated cells (U) and cells irradiated with 6 kGy at different post-irradiation time points (PIR) were visualized on the gel immediately after irradiation (0) and at specific incubation times (in hours). Lambda PFG molecular mass standards were loaded in lane-M for size reference. The data suggests that the *ΔddrB ΔdprA* mutant exhibits relatively improved DSB repair compared to the *ΔddrB* mutant, while the *ΔpprA ΔdprA* mutant shows weaker DSB repair compared to the *ΔpprA* mutant.

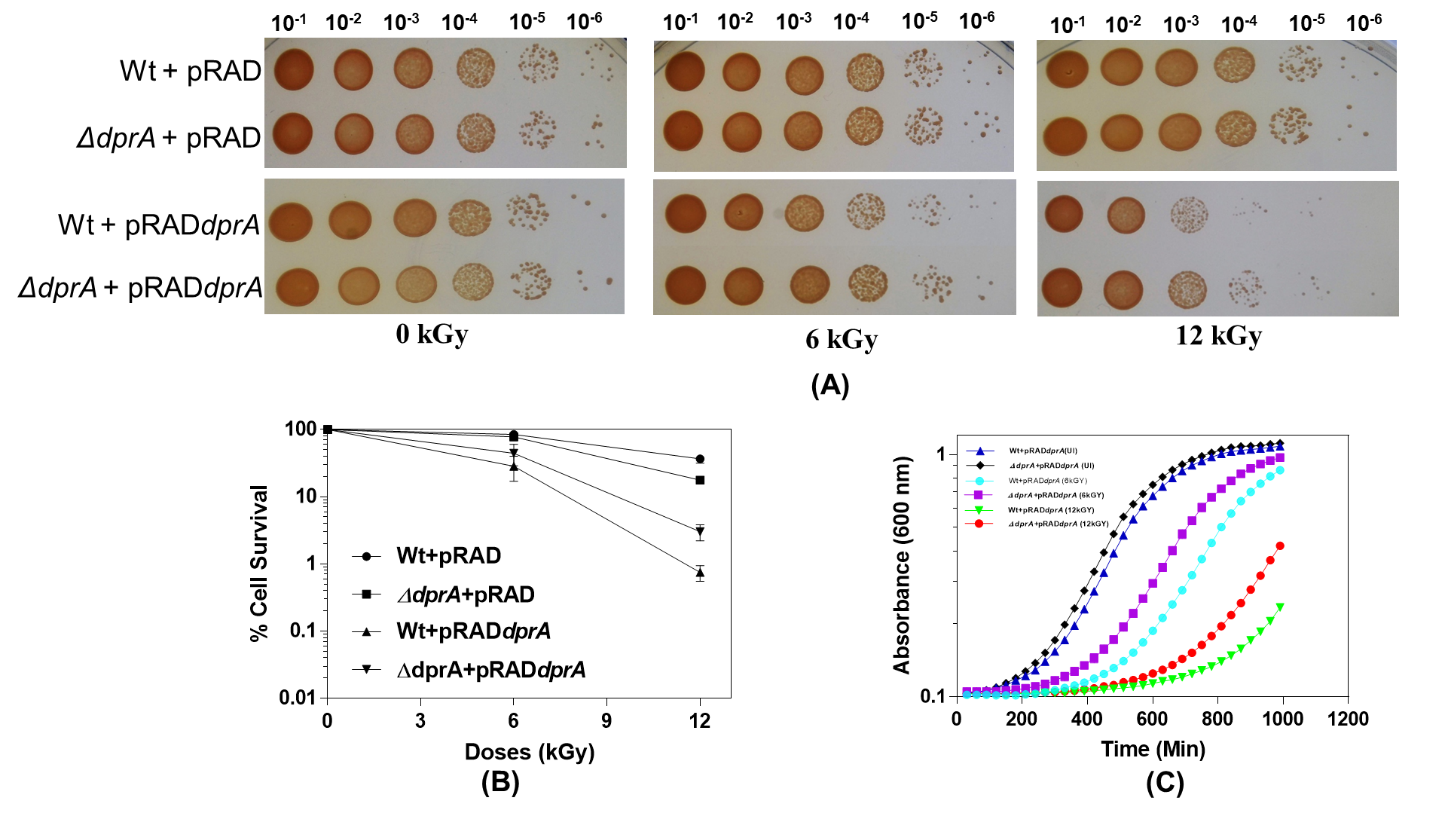

**Figure S7: The cell survival of wild type and *ΔdprA* mutant of *D. radiodurans* was assessed under gamma radiation by overexpressing DprA from the pRAD shuttle vector.** Both the wild type and *ΔdprA* mutant strains were transformed with either the pRAD control plasmid or the pRAD*dprA* plasmid for ectopic expression of DprA. (A) These transformed cells were exposed to various gamma radiation doses, and after dilution, aliquots were spotted on TGY plates and incubated at 32°C for 48 hours. (B) The percentage of cell survival fraction was plotted as a function of gamma radiation doses (kGy). (C) Additionally, the optical density at 600 nm was measured for the wild type and ΔdprA mutant strains overexpressing DprA under untreated conditions, as well as after exposure to 6 and 12 kGy of gamma radiation. The data suggested that ectopic overexpression of DprA negatively affects the cell survival and growth of *D. radiodurans*.

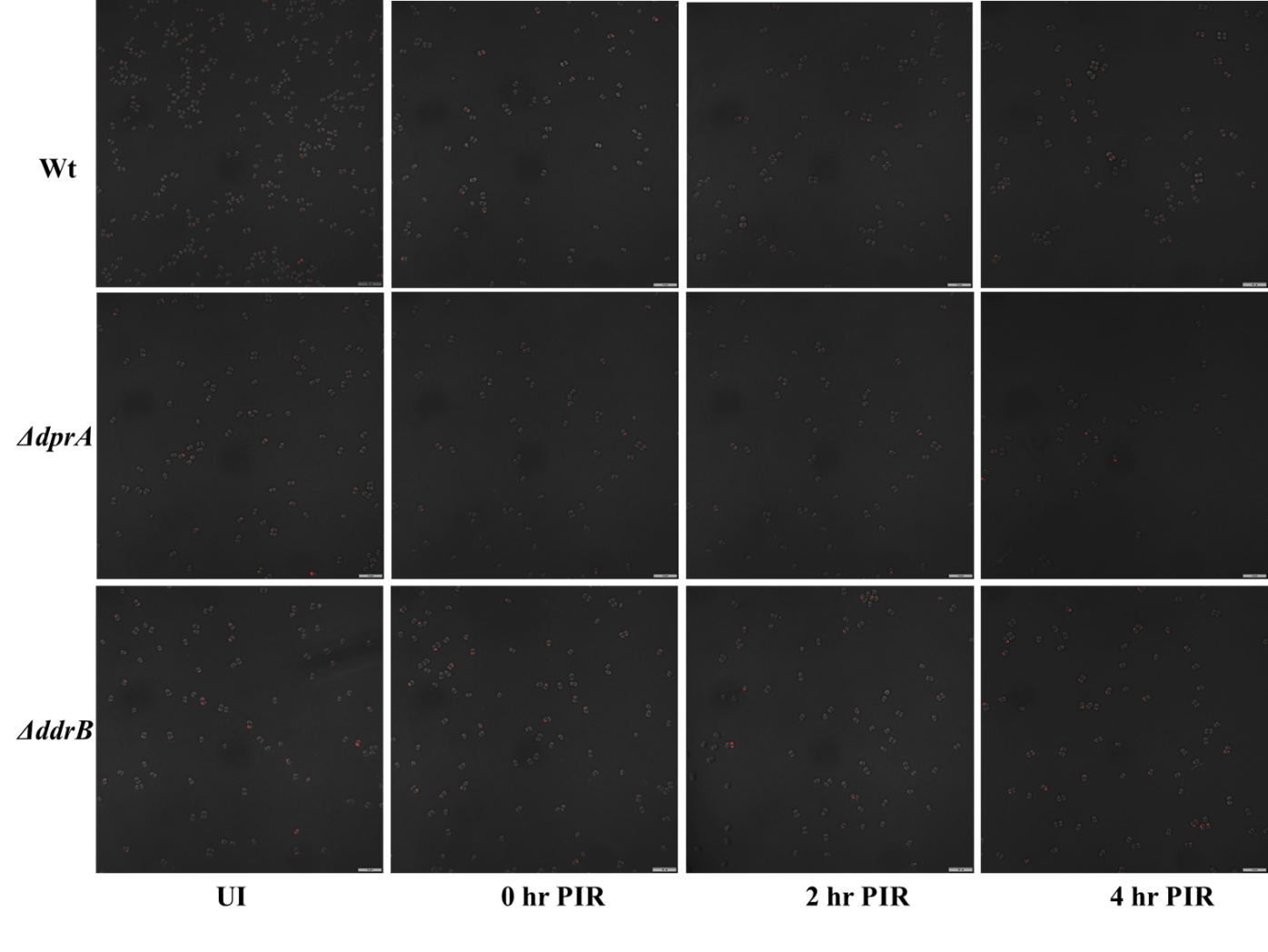

**Figure S8 RecA^RFP^ foci formation in wild type, *ΔdprA* and *ΔddrB* mutants.**

For fixed-cell imaging, *D. radiodurans* cells expressing pRADrecA-RFP were fixed and observed using Olympus FV3000 confocal microscope, specifically. These cells were examined in the RFP channel (561 nm) and under bright field (DIC) to determine the localization of RecA^RFP^. Image analysis was performed using automated cellSens software. The scale bar indicates a length of 10 µm.

**Table S1: Bacterial Strains, Plasmids, and Primers Used in This Study**

| **Bacterial strains** | | **Genotype** | **Source** |
| --- | --- | --- | --- |
| *D. radiodurans* R1 | | Wild type strain ATCC13939 | Lab stock |
| *E. coli* Novablue | | *end*A1 *hsd*R17*(r _K12_ ^−^ m _K12_ ^+^) sup*E44 *thi-1 rec*A1 *gyr*A96 *rel*A1 *lac*F’'*[pro*A^+^B*^+^ lac*I^q^ *Z∆*M15*::*T*n*10] (Tet^R^) | Lab stock |
| *∆pprA* | | *pprA Ω chl* | Prof. Issay Narumi lab |
| *ΔddrA* | | *ddrA Ω chl* | Prof. Pascale Servant lab |
| *ΔddrB* | | *ddrB Ω kan* | Prof. Pascale Servant lab |
| *ΔddrA ΔddrB* | | *ddrA Ω chl , ddrB Ω kan* | Prof. Pascale Servant lab |
| *ΔpprA ΔdprA* | | *pprA Ω chl, dprA Ω spec* | This work |
| *∆dprA* | | *dprA Ω spec* | This work |
| *ΔddrA ΔdprA* | | *ddrA Ω chl, dprA Ω spec* | This work |
| *ΔddrB ΔdprA* | | *ddrB Ω kan, dprA Ω spec* | This work |
| *ΔddrA ΔddrB ΔdprA* | | *ddrA Ω chl, ddrB Ω kan, dprA Ω spec* | This work |
| *D. radiodurans (thy^-^)* | | *thy^-^* | This work |
| *ΔdprA (thy^-^)* | | *dprA Ω spec, thy^-^* | This work |
| *ΔpprA (thy^-^)* | | *pprA Ω chl , thy^-^* | This work |
| *ΔpprA ΔdprA (thy^-^)* | | *pprA Ω chl, dprA Ω spec, thy^-^* | This work |
| *ΔddrB (thy^-^)* | | *ddrB Ω kan, thy^-^* | This work |
| *ΔddrB ΔdprA (thy^-^)* | | *ddrB Ω kan, dprA Ω spec, thy^-^* | This work |
| *ΔcomEA* | | *comEA (dr_1855) Ω kan* | This work |
| *ΔendA* | | *endA ( dr_1600) Ω kan* | This work |
| *ΔdprA* | | *dprA (dr_0120) Ω spec* | This work |
| *ΔpilT* | | *pilT (dr_1963) Ω spec* | This work |
| **Plasmids:** | | | |
| **Names** | **Characteristics and Source** | | lab stock |
| pVHS559 | A shuttle vector between *D. radiodurans* and *E. coli* (Spec^R^) | | lab stock |
| pRADgro | pRAD1 carrying 261bp *Bgl*II-*Xba*I fragment of promoter (Pgro) from *D. radiodurans* | | This work |
| pDSRED-*recA* | pDSRED carrying dr*recA* at *BamHI* and *KpnI* | | [64] |
| pRAD*recA*-RFP | pRAD carrying dr*recA* at *ApaI* and *EcoRV* | | [64] |
| pNOK*comEA* | pNOK suicidal vector carrying upstream and downstream DNA fragment of *comEA* gene (*dr_1855*) | | This work |
| pNOK*endA* | pNOK suicidal vector carrying upstream and downstream DNA fragment of *endA* gene (*dr_1600*) | | This work |
| pNOS*dprA* | pNOS suicidal vector carrying upstream and downstream DNA fragment of *dprA* gene (*dr_0120*) | | This work |
| pNOS*pilT* | pNOS suicidal vector carrying upstream and downstream DNA fragment of *pilT* gene (*dr_1963*) | | This work |
| **Primer details** | | | |
| **S. No.** | **Name of primer** | **Sequence (5’ to 3’)** | **Purpose** |
| 1 | *comEA*-UF | AAA GGG CCC GGC AAT CAC GTT CAT CAG | *ΔcomEA* mutant generation |
| 2 | *comEA*-UR | ATA GGA TCC GCG TGA CGC TCG CAG TGG |  |
| 3 | *comEA*-DF | ATA GGA TCC GAA GCC CTT CCC AAA GTC |  |
| 4 | *comEA*-DR | TAG TCT AGA AGA AGC CCT CAT CAA ACG |  |
| 7 | *endA*-UF | ATA GGT ACC TGC CCG ACT TCC TGC ATT | *ΔendA* mutant generation |
| 8 | *endA*-UR | AAA GGG CCC AGG AAA AGA GGT GGT GCA |  |
| 9 | *endA*-DF | ATA GGA TCC TTG CCC CTT AAT GCA GCA |  |
| 10 | *endA*-DR | TAG TCT AGA TAA CGG CGG GGT GAA ACG |  |
| 11 | *dprA*-UF | ATA GGT ACC TAA GCG CCT ATC AAG CCC TC | *ΔdprA* mutant generation |
| 12 | *dprA*-UR | AAA GGG ATG CCG CCG CAG GTT TTC GAT |  |
| 13 | *dprA*-DF | ATA GGA TCC GAA CTG AAC AAG GCG GCA GA |  |
| 14 | *dprA*-DR | TAG TCT AGA ACG CGG CAA CAG AGA GAA GTC |  |
| 15 | *pilT*-UF | AAA GGG CCC CAT CGA GAA CTT CAA CAT | *ΔpilT* mutant generation |
| 16 | *pilT*-UR | ATA GGA TCC ATG AAC TCG ATG GGG TCT T |  |
| 17 | *pilT*-DF | ATA GGA TCC AGC TCG CCA ACA ACC TCG T |  |
| 18 | *pilT*-DR | ATG TCT AGA AAC GAT TCC ACC GCA ATC GG |  |
